# Charting oncogenicity of genes and variants across lineages via multiplexed screens in teratomas

**DOI:** 10.1101/2021.03.09.434648

**Authors:** Udit Parekh, Daniella McDonald, Amir Dailamy, Yan Wu, Thekla Cordes, Kun Zhang, Ann Tipps, Christian Metallo, Prashant Mali

## Abstract

Deconstructing tissue-specific effects of genes and variants on proliferative advantage is critical to understanding cellular transformation and to systematic selection of cancer therapeutics. Dissecting these specificities at scale requires integrated methods for multiplexed genetic screens tracking fitness across time, across human cell types, and in a suitable cellular niche since functional differences also depend on physiological cues. Towards this, we present a novel approach, harnessing single-cell cancer driver screens in teratomas coupled with hit enrichment by serial teratoma reinjection, to simultaneously screen drivers across multiple lineages *in vivo*. Using this system, we analyzed population shifts and lineage-specific enrichment for 51 cancer associated genes and gene variants, profiling over 100,000 cells spanning over 20 lineages, across two rounds of serially injected teratomas. We confirmed that *c-MYC* alone or combined with myristoylated *AKT1* potently drives proliferation in progenitor neural lineages, demonstrating signatures of malignancy. These drivers directed teratoma development to lineages representative of pediatric tumors such as medulloblastoma and rhabdomyosarcoma. Additionally, mutant *MEK1^S218D/S222D^* provides a proliferative advantage in mesenchymal lineages like fibroblasts. Our method provides a powerful new platform for multi-lineage longitudinal study of oncogenesis.

## INTRODUCTION

The onset of cancer is often posited to be an evolutionary process enabled by the acquisition of somatic mutations and genetic lesions over time^1, 2^. Yet, these mutations lead to survival advantages and cancer only if accumulated in the relevant types of cells at the appropriate stage of differentiation, at the appropriate time and in the appropriate cell state^3–5^. Understanding this process of tumorigenesis and dissecting the tissue-specific molecular mechanisms which govern neoplastic transformation is a longstanding goal in cancer biology. In particular, with the explosion in cancer genome sequencing data^6–11^, understanding the tissue-specific oncogenicity of the growing list of genetic variants of unknown significance may lead to significant advances in building early detection systems, which would improve patient outcomes^12^, as well as inform the development and application of therapeutic strategies^13^.

In this regard, xenograft and genetically engineered animal models^14–16^ have been especially useful to recapitulate the process by which healthy cells undergo transformation, but such models often do not completely capture human biology, transformation or tumorigenesis^14–18^. On the other hand, immortalized cell lines and primary cells have been important workhorses of cancer research^19, 20^, aiding in mechanistic studies, the uncovering of therapeutic vulnerabilities and understanding resistance mechanisms^21^. Yet, elucidating the wide spectrum of tissue-specific programs governing transformation^22–30^ remains a challenge^31^, especially as access to and culture of diverse types of primary cells is challenging. Such *in vitro* systems also exclude the environmental and physiological context which are key modulators of driver-specificity, and often lack the ability to perturb cells along their differentiation trajectories or in distinct states, an important factor in driver-specific transformation^32^.

Systems which allow us to ethically gain access to developing human tissue models, recapitulating the architecture and signaling programs of native tissue, in a suitable physiological niche could be an invaluable tool to deconstruct tissue-specific drivers of neoplastic transformation. In this regard, there has been significant work toward the use of directed differentiation of stem cells^33^ in 2-dimensional monolayer culture as well as into organoids^34–41^ to model cancer. These models capture various cell states along the developmental trajectory, while organoids also capture the diversity of cells present in specific tissue, organized in native-like architecture. However, these systems are cultured in specialized conditions, which may not represent *in vivo* settings, and lack vasculature which is a central characteristic of cancers^42^. *In vivo* engraftment of suitably differentiated human stem cells provides an avenue to study the human-specific dynamics of oncogenesis and cancer evolution in an appropriate milieu^43–46^. But, each of these systems provides access to a single or a few lineages at a time, thus, to test drivers in a panoply of lineages to assess their oncogenic potential in each is a laborious and slow process.

Recently, our group demonstrated that human pluripotent stem cell (hPSC) derived teratomas, which are typically benign tumors with differentiated cells from all three germ layers and regions of organized tissue-like architecture^47^, can enable high throughput genetic screens simultaneously across cell types of all germ layers^48^. Leveraging this, we propose here a method that uses the teratoma model, in combination with single cell RNA-sequencing (scRNA-seq) based open reading frame (ORF) overexpression screens^49^ and serial tumor propagation, to massively parallely assess the tissue-specific oncogenicity of genes and gene variants across a diverse set of lineages.

## RESULTS

### Design of a multiplex *in vivo* screening platform

To enable a screening platform which would simultaneously allow the determination of lineages along with detection of perturbations, we implemented pooled overexpression screens in teratomas with scRNA-seq readout, using a lentiviral overexpression vector we previously developed for compatibility with scRNA-seq based pooled genetic screens^49^ (**Figure 1a**). The vector contains a unique 20 bp barcode for each library element, located 200 bp upstream of the lentiviral 3’ long terminal repeat (LTR). This yields a polyadenylated transcript with the barcode proximal to the 3’ end, thus allowing detection in droplet based scRNA-seq systems which rely on poly-A capture. Each cancer driver was cloned into the lentiviral backbone vector, packaged individually into lentivirus particles and this individually packaged lentivirus then combined, to avoid barcode shuffling due to the recombination driven template switching inherent in pooled lentiviral packaging^49–52^. To allow for combinatorial driver transduction, these driver libraries were then transduced into H1 human embryonic stem cells (hESCs), which we had previously characterized to be of normal karyotype^48^. The hESCs were transduced at the highest titer where we did not see visible morphological changes, and maintained under antibiotic selection for 4 days after transduction (**Figure 1b, Supplementary Figure S1, Methods**). In addition, for cells profiled prior to injection by scRNA-seq, for the majority of drivers we observed no significant changes per cluster in the proportion of cells expressing drivers as compared to those expressing the internal negative control (**Supplementary Figure S1**), confirming that the driver library was not significantly perturbing the hESCs from their basal state.

**Figure 1:**
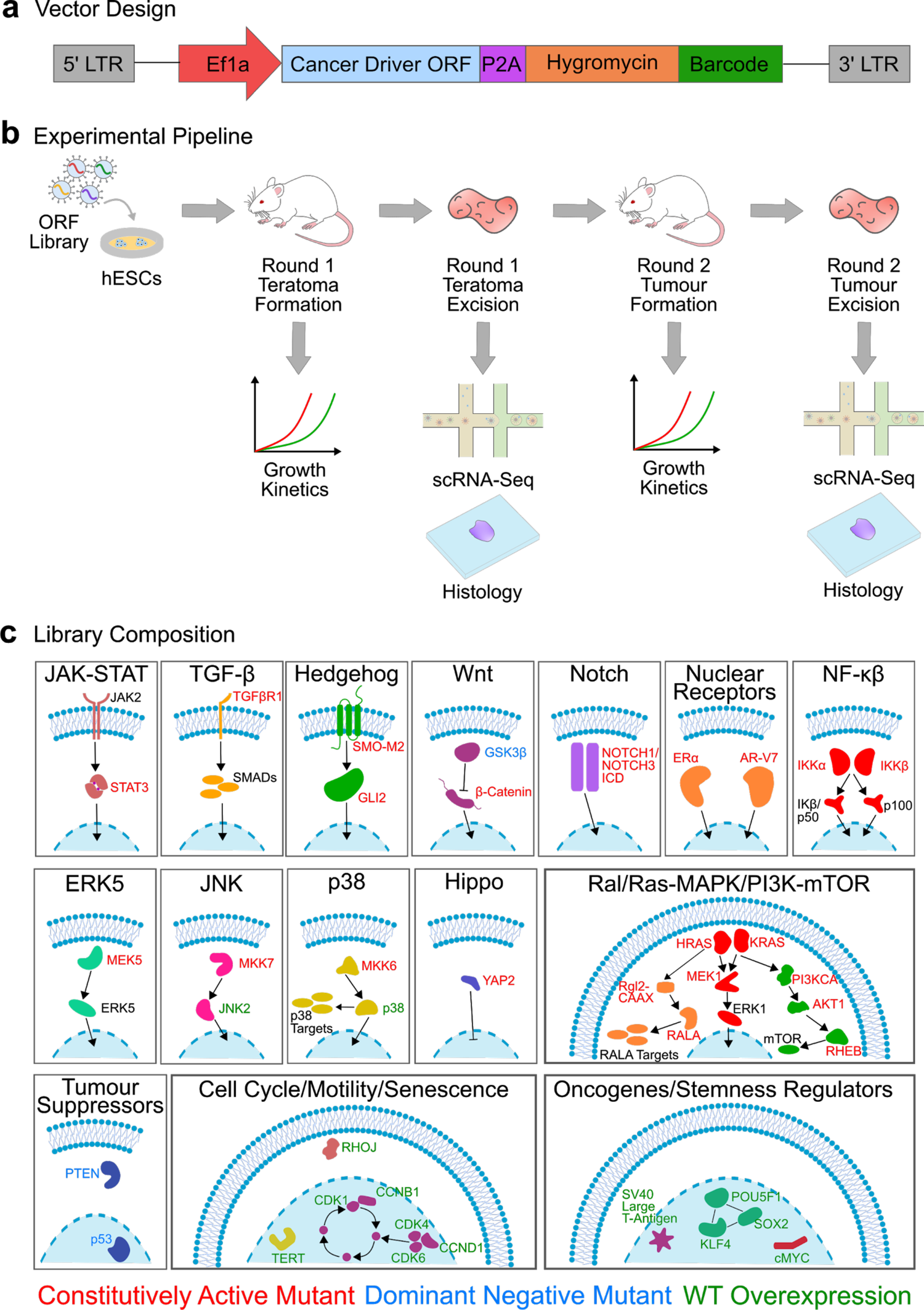
Overview of experimental design and library construction. (**a**) Schematic of lentiviral overexpression vector. (**b**) Schematic of experimental framework for evaluation of effects of cancer driver overexpression in developing teratomas and reinjected tumors: Individual cancer driver ORFs are cloned into the barcoded ORF overexpression vector, packaged into lentivirus then pooled for transduction of hESCs. Transduced cells are then injected subcutaneously into Rag2^-/-^;γc^-/-^ immunodeficient mice to form the round 1 teratomas. Teratoma growth is assayed by caliper-based measurement of approximate elliptical area of the tumor. Once the teratoma size reaches a threshold value, teratomas are excised and assayed by single-cell RNA-seq and histology, and a fraction of cells are reinjected in Rag2^-/-^;γc^-/-^ immunodeficient mice to form round 2 tumors for further enrichment of drivers. Round 2 tumor growth is also monitored via caliper-based measurements and at the end point tumors are assayed via scRNA-seq and histology. (**c**) Composition of the cancer driver ORF library, encompassing major signaling pathways, oncogenes and stemness regulators involved in oncogenesis and cancer progression.

To form teratomas with these library-transduced hESCs, we subcutaneously injected 6-8 million of these cells suspended in a 1:1 mixture of Matrigel and the pluripotent stem cell medium mTeSR1 in the right flank of anesthetized Rag2^-/-^;γc^-/-^ immunodeficient mice (**Figure 1b, Methods**). As growth controls, wild type, unmodified H1 hESCs were similarly injected in a separate group of Rag2^-/-^;γc^-/-^ immunodeficient mice. The growth of all teratomas was monitored via weekly caliper-based measurements of elliptical area (**Figure 1b, Methods**). Once the teratomas reached a threshold size for excision, they were extracted for downstream processing. After extraction, tumors were weighed and measured, and representative pieces were frozen for cryosectioning and histological analysis. The remaining pieces were dissociated into single cell suspensions. A part of this suspension was processed for single cell RNA-sequencing (scRNA-seq) using the droplet based 10X Genomics Chromium system, while a second part consisting of 6-8 million cells was reinjected into Rag2^-/-^;γc^-/-^ immunodeficient mice for serial proliferation to further enrich transformed lineages and drivers of proliferative advantage. These reinjected round 2 tumors were monitored and processed similarly to the initially formed round 1 teratomas (**Figure 1b, Methods**), to obtain histology information as well as transcriptomic profiles via scRNA-seq.

### Deconstructing fitness effects across lineages

Using this method, we screened 51 genes and gene variants representing major signaling pathways and oncogenes^21^ involved in oncogenesis, tumor proliferation, survival, cell cycle regulation and stemness regulation (**Figure 1c**).

To screen these drivers, we generated four teratomas from the perturbed hPSCs. In these first-round driver library teratomas, we observed palpable and measurable tumors between 20-27 days after injection, which grew to a size sufficient for extraction between 41-60 days after injection. In comparison, out of eight teratomas formed from unperturbed wild type cells, one did not form a tumor in 90 days of monitoring, while the seven remaining teratomas were palpable and measurable between 27-39 days after injection and six of them grew to a size sufficient for extraction between 55-78 days after injection (**Figure 2a**).

**Figure 2:**
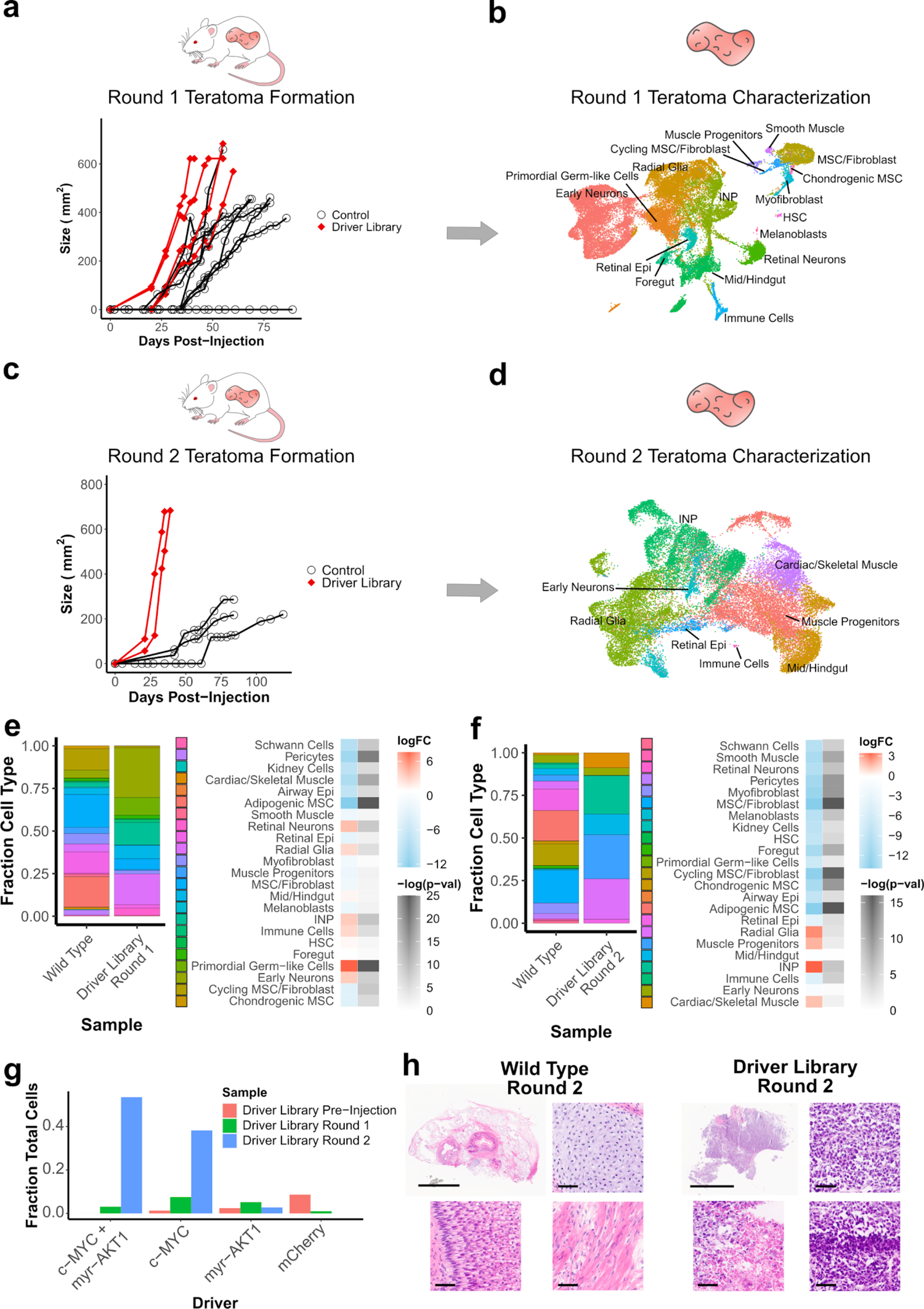
Identification of significantly enriched drivers and cell types. (**a**) Growth kinetics of round 1 teratoma formation for injections with driver library transduced hESCs vs control WT hESCs (**b**) UMAP visualization of cell types from round 1 teratomas formed by driver library transduced hESCs. (**c**) Growth kinetics of round 2 tumors formed from re-injected cells from round 1 teratomas formed by driver library transduced hESCs vs WT hESCs. Control measurements are from a common set of tumors grown from the parent WT hESC line, which were used as growth controls for all experiments in this study. (**d**) UMAP visualization of cell types from round 2 tumors formed by re-injected cells from round 1 teratomas of driver library transduced hESCs. (**e**) Relative fraction of each cell type in round 1 teratomas formed from library transduced hESCs and WT hESCs, and log fold change of each cell type for driver library teratomas vs WT teratomas. (**f**) Relative fraction of each cell type in round 2 tumors formed from library transduced hESCs and WT hESCs, and log fold change of each cell type for driver library tumors vs WT tumors. (**g**) Relative fraction of top enriched drivers prior to injection and in each round of tumor formation. (**h**) H&E stained sections of round 2 tumors formed from WT and driver library transduced hESCs. WT tumors display mature cell types from all three germ layers, such as cartilage (top right), muscle (bottom right) and dermis-like epithelium (bottom left). Driver library tumors display disorganized and more homogenous composition along with markers of transformation such as nuclear pleiomorphism (top right), areas of high mitotic rates (bottom right) and areas of necrosis (bottom left). Scale bars for full section are 5 mm, scale bars for magnified images are 50 μm.

Once these round 1 driver library teratomas were excised and processed via scRNA-seq, gene expression matrices were generated from the resultant data using 10X Genomics cellranger. We then used these expression matrices with the Seurat pipeline^53^ to integrate data from all four driver library teratomas and compensate for batch effects. Using this integrated data matrix, cells were then clustered using a shared nearest neighbor algorithm. Cell types were classified by the Seurat label transfer process using a previously classified set of teratomas generated from wild type H1 hESCs^48^ as a reference. Clusters were projected as a uniform manifold approximation and projection (UMAP) scatterplot (**Figure 2b, Methods**), with cell types well distributed across all the teratomas (**Supplementary Figure S2a**). 17 out of 23 cell types detected in wild type teratomas were detected in the driver library teratomas.

Drivers were assigned to individual cells by amplifying paired scRNA-seq cell barcodes and library barcodes from the unfragmented scRNA-seq cDNA, enabling genotyping of each cell^49^ (**Methods**). Barcodes were detectably expressed in nearly 45% of cells in this round, and we hypothesize that stochastic silencing of the lentiviral cassette and potentially sparse capture during scRNA-seq may be leading to barcode association for a fraction of the captured cells. 54% of these genotyped cells expressed a single detectable barcode while the remaining genotyped cells expressed two or more barcodes (**Supplementary Figure S2b-d**).

A subset of the dissociated cells from each of the round 1 teratomas were reinjected to grow round 2 tumors (**Figure 1b, Methods**), to further enrich dominant drivers and lineages. From these four reinjected tumors, two tumors were processed via scRNAseq and histology. Three of the fastest growing wild type teratomas were also dissociated into single cell suspensions and reinjected to form round 2 control tumors.

For the two driver library tumors which were processed, tumors were palpable and measurable by 21 days after injection and reached the size threshold for excision by 35-39 days after injection. This was a markedly faster growth rate than the control round 2 tumors, which in contrast reached a detectable and measurable size at least 42 days after injection and did not reach the size threshold for excision in up to 120 days of monitoring (**Figure 2c**). Due to the hardness and density of the tissue, none of the control round 2 tumors could be dissociated into single cell suspensions using the standard dissociation protocols.

Post-excision, the round 2 driver library tumors were processed in a manner similar to the round 1 teratomas. Mapping clusters to cell types in wild type teratomas revealed only 7 out of the 23 cell types detected in the wild type teratomas, a sign of potential fitness advantage in these lineages leading to them dominating the tumor composition (**Figure 2d**). A significant difference was observed between the two tumor samples even after integrating the data with batch correction via the Seurat integration pipeline (**Supplementary Figure S2e**). This challenge in integrating scRNA-seq datasets is also observed in clinical tumor samples, which display patient- or tumor-specific batch effects^54–56^ due to inter- and intra-tumor heterogeneity. Barcodes were detectably expressed in 95% of cells in this round (**Figure S2e, Supplementary Figure S2f**), suggesting that cells with a survival and proliferation advantage were those where the lentiviral cassette was not silenced and was robustly expressed.

We then examined the cell type populations which were present in these driver library tumors, compared to teratomas derived from wild type hESCs. In the round 1 teratomas, immature neural lineages and neural progenitors dominated the composition of the tumor with early neurons, radial glia and intermediate neural progenitors (INPs) making up 60% of the teratoma, and primordial germ-like cells another 10% (**Figure 2e**). Compared to the control teratomas this represented a 4-6 fold increase in proportion of the neural cell types, and a nearly 100 fold increase in proportion of the primordial germ-like cells. In contrast, the mesenchymal lineages were significantly reduced as a proportion of cells in the tumors (**Figure 2e**). In the round 2 tumors, cell type populations were further shifted. The majority of these round 2 tumors consisted of neural progenitor-like cells, primarily radial glia and INPs, which made up 45% of these tumors, and muscle progenitor-like cells, which made up 25% of the tumors (**Figure 2f**). Here we observed significant sample-specific effects with tumor 1 composed of a majority of neural-like cells, while tumor 2 was composed primarily of muscle, muscle progenitor-like and gut-like lineages (**Supplementary Figure S2e, g**).

There was a dramatic redistribution of driver populations detected over the course of these serial injections. Prior to injection, *c-MYC* and myristoylated *AKT1* (*myr-AKT1*) constituted 1.3% and 2.4% respectively of total cells while a combinatorial transduction of the two, *c-MYC* + *myr-AKT1*, was undetectable in 5949 genotyped cells. In the first round of teratomas *c-MYC*, *myr-AKT1* and *c-MYC* + *myr-AKT1* constituted 16.7%, 11.6% and 6.9% of cells respectively, while strikingly in the second round of tumors *c-MYC* + *myr-AKT1* and *c-MYC* made up 56% and 40% of cells respectively (**Figure 2g**). These observations are consistent with the observation of elevated expression or amplification of *c-MYC*, or *c-MYC* dependent survival and proliferation, in subsets of embryonal tumors such as medulloblastoma^45, 46, 57, 58^ and atypical rhabdoid/teratoid tumors^59, 60^, and pediatric soft tissue tumors like rhabdomyosarcoma^61–63^. The observed enhanced combinatorial effect of the drivers is also in line with the cooperative action of *c-MYC* and *AKT1*, where the action of *c-MYC* is negatively regulated by the *FOXO* group of transcription factors, which in turn are regulated by the *PI3K/AKT* pathway^64^. A constitutively active form of *AKT1*, such as *myr-AKT1*, phosphorylates *FOXO* transcription factors and abrogates their function, thus allowing for uninhibited *c-MYC* activity^46, 64^. In our observations with the teratoma based system, *c-MYC* seemed to also drive a muscle-like phenotype which was reduced as a fraction of the population when *myr-AKT1* was expressed in combination. This may also explain the sample-specific clustering observed between the two tumors which had different drivers as the top enriched hits leading to fitness advantages (**Supplementary Figure S2h, i**).

Histology displayed clear differences between the control and driver library tumors. While control tumors had a diversity of cell types from all three germ layers in well-organized architectures (**Figure 2h**), the driver library tumors had far more homogenous composition and displayed distinct signs of malignancy such as nuclear pleiomorphism, areas of high mitotic rate and areas of necrosis (**Figure 2h**), suggesting the de novo creation of a transformed phenotype driven by *c-MYC* overexpression with or without the accompanying dysregulation of *AKT1* signaling.

### Screening less-dominant drivers by removing dominant hits

In the previously described screens, the proliferative advantage conferred by the top hits caused cells expressing those drivers to overwhelm all other cells, such that other drivers were not significantly detected in the second round of tumors formed by serial reinjection. To assay the fitness effects of other drivers, we hypothesized that removing the top hits from the library would allow the detection of enrichment of other drivers.

Towards this we repackaged a lentiviral driver sub-library, removing *c-MYC* and *myr-AKT1* from the pool. Using this sub-library we conducted the screening process similarly to that for the full driver library. We injected and monitored six round 1 teratomas, out of which we characterized the three fastest growing ones via scRNA-seq. Similar to before, round 1 teratomas were reinjected for driver enrichment, and tumors were monitored for 75 days, with the two fastest growing tumors assayed via scRNA-seq.

The round 1 driver sub-library teratomas were measurable between 18-25 days after injection and reached a size sufficient for extraction between 40-60 days after injection (**Figure 3a**). We again assessed the cell type populations in these sub-library screens by mapping the cells to the wild type teratomas as a reference. Using this, we determined that the round 1 teratomas contained 13 major cell types, a majority of which were neural cell types, in particular early neurons, radial glia and INPs, distributed across all three samples (**Figure 3b, Supplementary Figure S3a**). In these samples, barcodes were detectable in 27% of cells (**Supplementary Figure S3b-d**).

**Figure 3:**
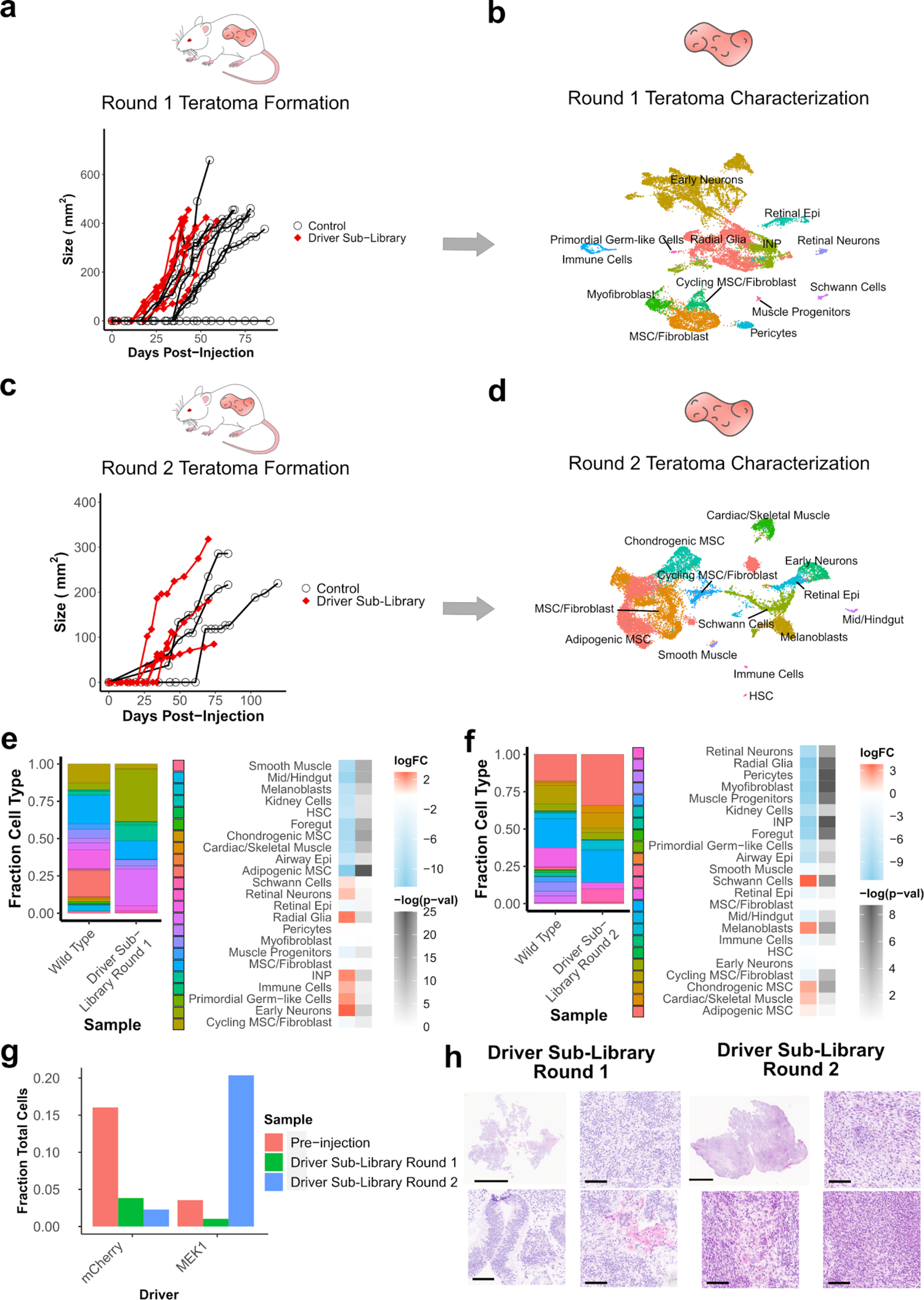
Identification of significantly enriched drivers and cell types for tumors formed with driver libraries without *c-MYC* and *myr-AKT1*. (**a**) Growth kinetics of round 1 teratoma formation for injections with driver library transduced hESCs vs WT hESCs (**b**) UMAP visualization of cell types from round 1 teratomas formed by driver library transduced hESCs. (**c**) Growth kinetics of round 2 tumors formed from re-injected cells from round 1 teratomas formed by driver library transduced hESCs vs WT hESCs. Control measurements are from the common set of tumors grown from the parent WT hESC line, which were used as growth controls for all experiments in this study. (**d**) UMAP visualization of cell types from round 2 tumors formed by re-injected cells from round 1 teratomas of driver library transduced hESCs. (**e**) Relative fraction of each cell type in round 1 teratomas formed from library transduced hESCs and WT hESCs, and log fold change of each cell type for driver library teratomas vs WT teratomas. (**f**) Relative fraction of each cell type in round 2 tumors formed from library transduced hESCs and WT hESCs, and log fold change of each cell type for driver library tumors vs WT tumors. (**g**) Relative fraction of top enriched drivers prior to injection and in each round of tumor formation. (**h**) H&E stained sections of round 1 and round 2 tumors formed from hESCs transduced with driver libraries without *c-MYC* or *myr-AKT1*. Round 1 tumors contain neuroectodermal and epithelial cell types as the majority of cells, while round 2 tumors contain mesenchymal cell types as the majority of cells. Scale bars for full section are 5 mm, scale bars for magnified images are 100 μm.

In contrast to the full driver library, the round 2 driver sub-library teratomas showed a slower growth rate, reaching a measurable size between 27-38 days after injection. While one of the round 2 tumors reached a sufficient size for extraction at 75 days after injection (**Figure 3c**), with the fastest growing tumors extracted and assayed via scRNA-seq. In these round 2 sub-library tumors, we again detected 13 major cell types, but as opposed to neural or muscle progenitor lineages, the majority were mesenchymal cell types with adipogenic mesenchymal stromal cells (MSCs), MSC/fibroblasts and chondrogenic MSCs (**Figure 3d, Supplementary Figure S3e**). Barcodes were detectable in 46% of cells in this round of tumors (**Supplementary Figure S3f**), with the increase in the fraction of genotyped cells suggesting that surviving cells which express barcodes may have a fitness or survival advantage conferred by the expressed driver. We again subsetted the drivers to visualize only those detected in 25 cells or more (**Supplementary Figure S3g, h**), the majority of which were driven wholly or in part by *MEK1^S218D/S222D^*, while the remaining were driven by *RHOJ*, a small GTP-binding protein known to play a role in cell migration, which has recently been found to confer proliferative advantage in multiple lineages^31^.

We then evaluated the cell type populations in comparison to teratomas formed from wild type H1 hESCs. In the round 1 sub-library teratomas, immature and progenitor neural cell types constituted 70% of the total population of the teratoma, and fibroblast-like mesenchymal lineages making up a further 20% of the teratoma (**Figure 3e**). However, in the round 2 tumors, in a striking difference compared to the full library screens, the tumors were composed primarily of fibroblast and MSC phenotypes, which constituted 65% of the tumors (**Figure 3f**). Along with the fibroblast lineages which constituted a majority of these tumors, a small neural component persisted via an expansion in Schwann cells and melanoblasts, which may be derived from Schwann cell precursors^65^. The neural progenitor-like lineages present in the *c-MYC* and *myr-AKT1* driven tumors were not present in these tumors. This change in composition of the tumors was accompanied by an enrichment of the constitutively active mutant *MEK1^S218D/S222D^* which was present in 3.3% of cells prior to injection but constituted 20% of all cells in the round 2 tumors. In comparison, the fraction of cells expressing the internal negative control, mCherry, fell from 16% of cells prior to injection to 2% of cells in the round 2 tumors (**Figure 3g**). This role of *MEK1^S218D/S222D^* in supporting proliferation and survival of fibroblasts is consistent with previously reported results in literature where expression of constitutively active versions of *MEK1* were sufficient to trigger proliferative states and even transformation in fibroblasts *in vitro*^66–68^.

Histology images from the two rounds of tumors further confirmed the cellular composition. In round 1 teratomas we observed a majority of neuroectoderm-like cell types, while in round 2 tumors the majority of cells were mesenchymal in nature (**Figure 3h**). In these sub-library screens, we did not observe clear indications of malignancy as we observed in the *c-MYC* and *myr-AKT1* driven full library screens, suggesting that while *MEK1^S218D/S222D^* may drive proliferative advantage, it was a less potent driver of cellular transformation on its own.

### Validating enriched driver hits

To validate the hits obtained from the two sets of screens we individually tested the effects of *c-MYC*, *myr-AKT1*, *c-MYC + myr-AKT1* and *MEK1^S218D/S222D^*. hPSCs were transduced with either one of these driver vectors or a negative control vector expressing mCherry. Prior to injection the driver and control transduced cells were mixed in a 1:1 ratio, with a portion of these mixed cells pelleted and genomic DNA extracted to assess barcode distribution, and the remaining cells injected for teratoma formation in three Rag2^-/-^;γc^-/-^ immunodeficient mice for each driver to be validated. Teratomas were formed from these injected cells, excised when they reached sufficient size, representative pieces of the excised tumors were immediately flash frozen in liquid nitrogen and some representative pieces preserved for cryosectioning, while the remaining tissue was dissociated. Dissociated cells were divided to be stored for genomic DNA extraction, RNA extraction and for serial reinjection to form round 2 tumors. Round 2 tumors were also then allowed to grow to a size sufficient for extraction and then excised and dissociated as for round 1. Dissociated cells were stored for genomic DNA extraction and RNA extraction.

Consistent with the observations during the screens, teratomas formed by hESCs transduced with *c-MYC* and *c-MYC + myr-AKT1* grew at the highest rate compared to all other tumors. In the round 1 tumors, the *c-MYC*, *myr-AKT1* and *c-MYC + myr-AKT1* were measurable between 27-32 days after injection and were at an extractable size between 42-48 days after injection. While for *MEK1^S218D/S222D^* round 1 teratomas, tumors were measurable 33-40 days after injection and at an extractable size 59-78 days after injection (**Figure 4a**). In the round 2 tumors, those driven by *c-MYC + myr-AKT1* grew at the fastest rate, followed by *c-MYC*, both of which showed a significantly enhanced growth rate compared to the control round 2 tumors. Round 2 tumors driven by *myr-AKT1* alone or those driven by *MEK1^S218D/S222D^* did not show a significantly enhanced growth rate compared to the control tumors (**Figure 4a**), although tumors driven by *MEK1^S218D/S222D^* clustered toward the higher end of the control tumor growth rate.

**Figure 4:**
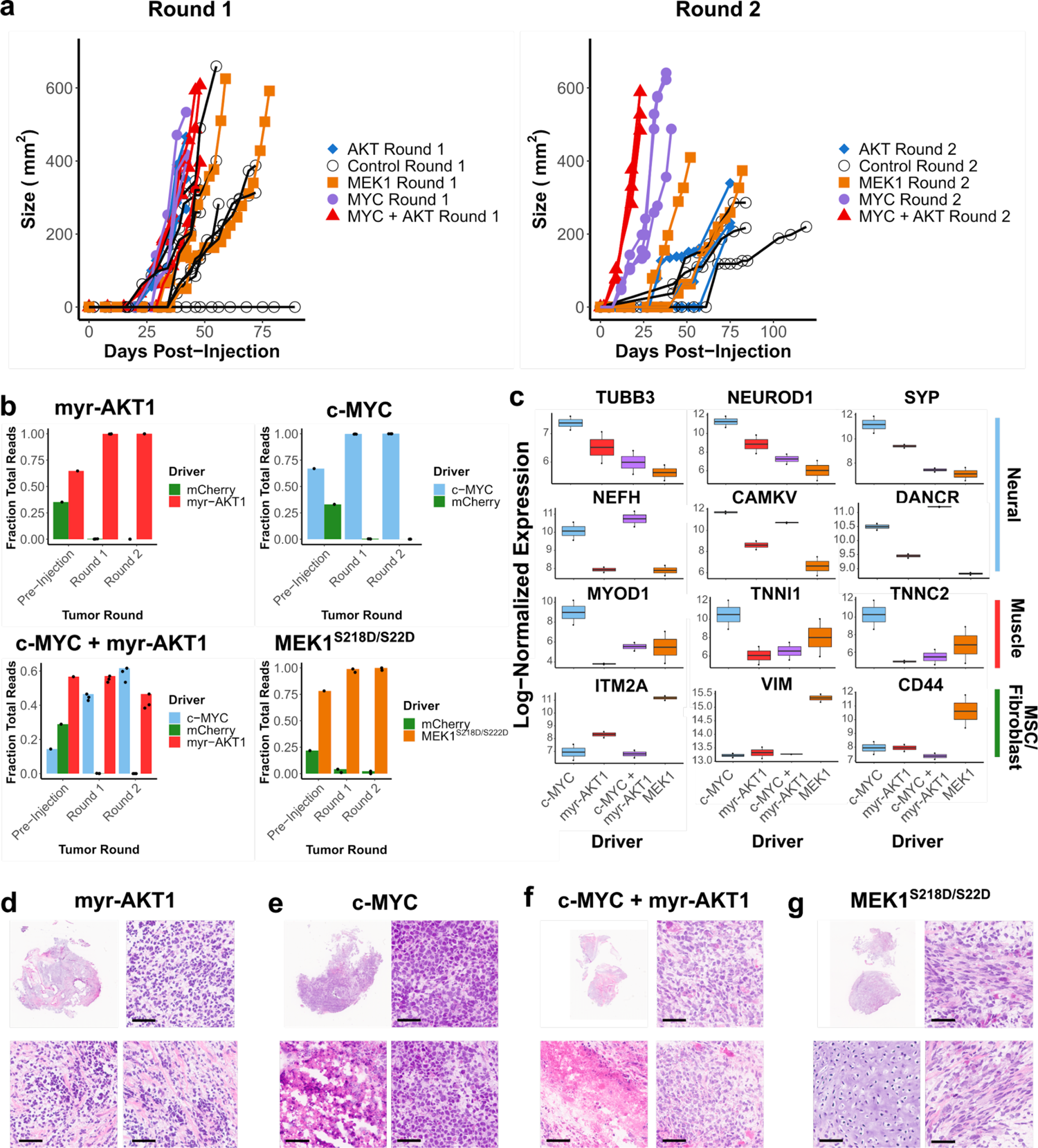
Validation of tumor formation and proliferative advantages of significantly enriched drivers from library screens. (**a**) Growth kinetics of round 1 and round 2 tumors formed from a mixture of hESCs transduced with driver hits (c-MYC, myr-AKT1, c-MYC + myr-AKT1 or MEK1 (S218D, S222D) and hESCs transduced with a negative control (mCherry). Control measurements are from the common set of tumors grown from the parent WT hESC line, which were used as growth controls for all experiments in this study. (**b**) Fraction of reads detecting either the driver or negative control barcodes at each stage: pre-injection, round 1 tumor formation and round 2 tumor formation. Barcodes were amplified from genomic DNA. (**c**) Gene expression of lineage-specific markers for round 2 tumors driven by individual hits. Expression values are normalized and log transformed. (**d**) H&E stain of round 2 tumor driven by myr-AKT1 showing regions of poor differentiation (top right) and necrosis (bottom right) interspersed with regions of organized tissue (bottom left). (**e**) H&E stain of round 2 tumor driven by c-MYC showing regions of poor differentiation (top and bottom right) and regions of necrosis (bottom left). (**f**) H&E stain of round 2 tumor driven by c-MYC + myr-AKT1 showing regions of poor differentiation (top and bottom right) and regions of necrosis (bottom left). (**g**) H&E stain of round 2 tumor driven by MEK1 (S218D, S222D) showing primarily regions of mature mesenchymal fibroblast-like tissue (top and bottom right) and cartilage-like (bottom left). Scale bars for magnified images are 50 μm.

To assess the barcode distribution from cells prior to injection through both rounds of tumor formation, we PCR amplified barcodes from genomic DNA isolated from stored cell pellets and quantified them via deep sequencing. We find that negative control barcodes decrease from 21-35% of total mapped reads prior to injection, to less than 2% of reads for *MEK1^S218D/S222D^*, and less than 0.1% of reads for all other drivers by the second round of tumors (**Figure 4b**).

We then performed bulk RNA-sequencing on these validation tumors to assess their composition. In agreement with the results of the screens, we found that the *c-MYC* driven tumors were composed of a mix of neural and muscle lineages displaying elevated expression of neural-related genes *TUBB3*, *NEUROD1*, *SYP*, *CAMKV* and *NEFH* (**Figure 4c**). In comparison, tumors driven by a combination of *c-MYC* and *myr-AKT1* were primarily composed of a neural progenitor like phenotype, expressing *NEFH* and *CAMKV* and elevated levels of *DANCR*, a non-coding RNA which acts to suppress differentiation and is present in many cancers^69, 70^ (**Figure 4c**). On the other hand, tumors driven by *MEK1^S218D/S222D^* displayed elevated expression of mesenchymal markers *VIM*, *ITM2A* and *CD44* (**Figure 4c**), which is again consistent with the results from the sub-library screens. We also compared the *c-MYC* and *MEK1^S218D/S222D^* driven tumors to pediatric cancers from the TARGET initiative. Using the 2000 most variable genes across the TARGET tumors we performed a principal component analysis (PCA), (**Supplementary Fig 4a**). Plotting the two PCs capturing the majority of variation, we observed that *c-MYC* driven tumors clustered toward the TARGET neuroblastoma tumors, while those driven by *MEK1^S218D/S222D^* clustered toward the kidney tumors (Wilm’s tumor and clear cell sarcoma of the kidney) and osteosarcoma.

*c-MYC* and *myr-AKT1* also contribute to metabolic reprogramming in tumors by promoting nucleotide biosynthesis^71, 72^. To further characterize *c-MYC* and *c-MYC* + *myr-AKT1* driven tumors, we quantified nucleobase levels via mass spectrometry. Compared to WT teratomas, *c-MYC* and *c-MYC* + *myr-AKT1* driven tumors increased the abundance of purine nucleobases, especially guanine (**Supplementary Figure 4b, d**). Further, this was supported by broad enrichment of nucleobase synthesis terms, and particularly purine synthesis related Gene Ontology terms in genes which were upregulated compared to WT teratomas (**Supplementary Figure 4c, e**). This is consistent with previous studies which have demonstrated the regulation of nucleotide metabolism by *c-MYC*^71, 73^, with de novo purine synthesis especially implicated in tumor maintenance^74^ and response to therapy^75^.

Histology showed that round 2 tumors driven by *myr-AKT1* alone showed poorly differentiated neural lineage cells interspersed with mesenchyme (**Figure 4d**). Round 2 tumors driven by *c-MYC* (**Figure 4e**) and those driven by *c-MYC + myr-AKT1* (**Figure 4f**) were composed of poorly differentiated cells with signs of malignancy, including necrosis. Finally, as observed in screen results, round 2 tumors driven by *MEK1^S218D/S222D^* were composed primarily of mesenchymal, fibroblast-like cells interspersed with cartilage (**Figure 4g**). These validation results strongly support the observations from the multiplexed screens and confirm the significant fitness advantage conferred by *c-MYC* and *c-MYC + myr-AKT1* on neural lineages, and of *MEK1^S218D/S222D^* on fibroblasts.

## DISCUSSION

To study the important problem of what drives oncogenic transformation in human tissue, investigators have relied on *in vitro* and animal model systems. While immense progress has been made using these, limitations still remain. Animal models retain significant differences with human biology, while *in vitro* systems lack vasculature and the physiological cues present in the *in vivo* niche which are involved in regulating lineage-specific transformation. Additionally, currently prevalent models often necessitate the investigation of a single or a few lineages at a time, raising an impediment in studying tissue-specificity across multiple cell types.

In this study, we have developed a novel platform to study the effects of cancer drivers across lineages which harnesses hPSC-derived teratomas to access a diverse set of lineages, ORF overexpression libraries to express cancer drivers, scRNA-seq to read out transcriptomic profiles which determine the cell type and detect the perturbing driver, and serial proliferation of the tumors to enrich drivers and lineages enjoying fitness advantages. Using this platform, we initially screened key cancer drivers across more than 20 cell types and de novo generated a transformed phenotype in a neural lineage via the overexpression of *c-MYC* and *myr-AKT1*, a constitutively active form of *AKT1*, which dominated the serially reinjected tumors and displayed the hallmarks of malignancy. To screen less-dominant drivers, we repeated the screens with *c-MYC* and *myr-AKT1* removed from the library, and captured the proliferative advantage of other drivers, such as the one conferred on mesenchymal lineages like fibroblasts via the overexpression of constitutively active *MEK1^S218D/S222D^*. These results were then validated by individually overexpressing these top hits during teratoma formation and serial reinjection, to confirm the enrichment of lineages detected in the screens.

While offering a powerful new approach in studying transformation, certain limitations and challenges merit consideration both in terms of interpretation of the resulting data, as well as in design of future implementations of the same. Firstly, in its current form, the overexpression vector is designed to be constitutively on. This may confound results since some drivers and lineages could be enriched due to their biased differentiation or inability to escape a state being governed by driver expression itself. In future versions of this platform, that effect may be mitigated by using an inducible expression system or one with recombinase-based control, both of which can be controllably turned on at specific time points, to ensure differentiation is not affected by driver expression. Additionally, background mutations and genetic alterations already present in hPSC lines may bias transformation phenotypes. Secondly, the predominantly embryonic versus adult state of the cell types might bias their innate transformation potential. In particular, we anticipate, due to the embryonic origin of the starting cells, this system might be especially applicable to modeling of pediatric tumors. Thirdly, the differentiation of teratomas is a partially random stochastic process, which leads to an inherent heterogeneity in the cell types available in each sample. To overcome this, cell fate engineering methods^48, 49^ may be paired with the screening platform to repeatably and predictably derive specific lineages of interest. Fourthly, while largely less discussed, dissociation and sample processing methods have a large impact on scRNA-seq results^76–78^. Improved protocols may allow for more fine-grained dissection of transcriptomes to assess cell state shifts within lineages. Fifth, the xenografted mouse models used for teratoma formation may be improved upon in two ways. In this study, we used a sub-cutaneous site of injection for teratoma formation, but the site of injection impacts teratoma differentiation^79^ as well as the lineage-specificity of cancer drivers. Orthotopic injections for specific tissues may provide a more appropriate niche. In addition, the mice used in this study are immune deficient, thus precluding the study of any effects of immune system interaction. A possible route to addressing this may be through the use of humanized mouse models^80^. Sixth, while in this demonstration we use scRNA-seq to determine cell type and state, epigenetic factors such as chromatin state are critical to tumorigenesis, cancer evolution and progression. Combining techniques to map epigenetic characteristics, such as ATAC-seq, may lead to a more detailed understanding of the determinants of tumor formation. Seventh, our pooled screening method may lead to transcriptomic and cell state shifts due to cell-cell communication and paracrine signaling. These challenges may potentially be addressed by utilizing newly developed spatial transcriptomics^81–83^ methods to tease apart cell endogenous versus exogenous effects. And finally, we have focused here primarily on deciphering the role of individual cancer drivers on oncogenic potential across lineages. Exploring combinatorial perturbations, especially in the background of tumor suppressor mutations will be crucial to dissecting both tumorigenicity as observed in native tumors, and also systematically studying variants-of-unknown significance, many of which individually may have only subtle phenotypes.

Taken together, we have demonstrated a proof of concept for a scalable, versatile platform which can screen multiple lineages and drivers in a single experiment, with a rich transcriptomic readout, thus providing a systematic path to studying the determinants and tissue-specificity of neoplastic transformation in human cells. We envision that refinements to this platform, coupled with the expanding array of available omics technologies will enable the comprehensive characterization of the trajectory of cells from normal to malignant states.

## Acknowledgements

This work was generously supported by UCSD Institutional Funds and NIH grants (R01HG009285, RO1CA222826, RO1GM123313). This publication includes data generated at the UC San Diego IGM Genomics Center utilizing an Illumina NovaSeq 6000 that was purchased with funding from a National Institutes of Health SIG grant (#S10 OD026929).

## Competing financial interests

P.M. is a scientific co-founder of Shape Therapeutics, Boundless Biosciences, Seven Therapeutics, Navega Therapeutics, and Engine Biosciences, which have no commercial interests related to this study. The terms of these arrangements have been reviewed and approved by the University of California, San Diego in accordance with its conflict of interest policies.

---

**Supplementary Figure 1:**
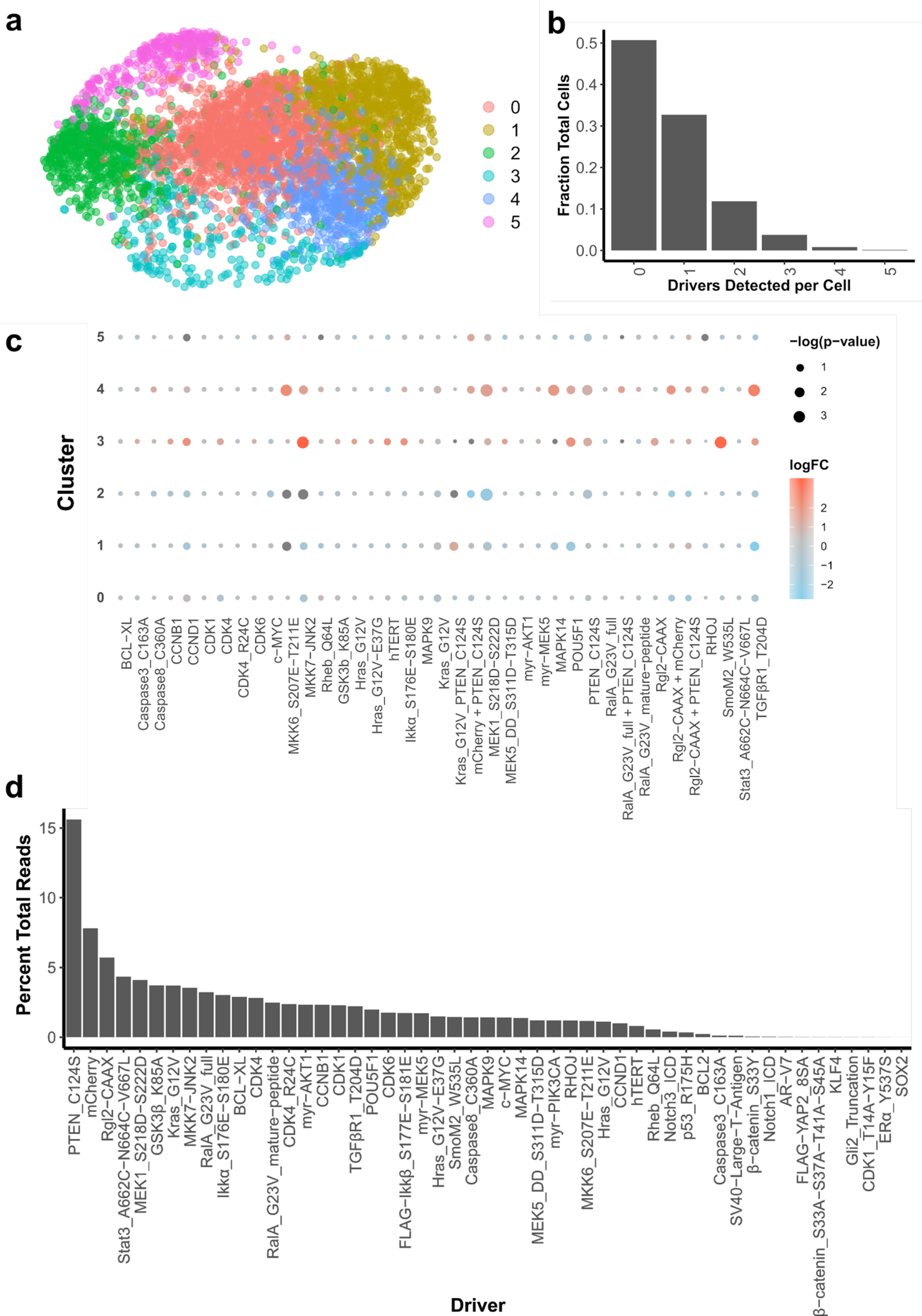
Characterization of driver library transduced cells prior to injection. (**a**) UMAP visualization of cells labelled by cluster identity. (**b**) Fraction of cells for each number of drivers detected in scRNA-seq for driver library transduced H1 hESCs. (**c**) Log2 fold change and significance of proportion of cells in a cluster expressing a driver versus proportion expressing the internal control (mCherry), calculated by permutation testing. Only drivers expressed in 20 or more cells are included. (**d**) Percentage of reads detecting each library element for barcodes amplified from genomic DNA extracted from driver-library transduced H1 hESCs prior to injection.

**Supplementary Figure 2:**
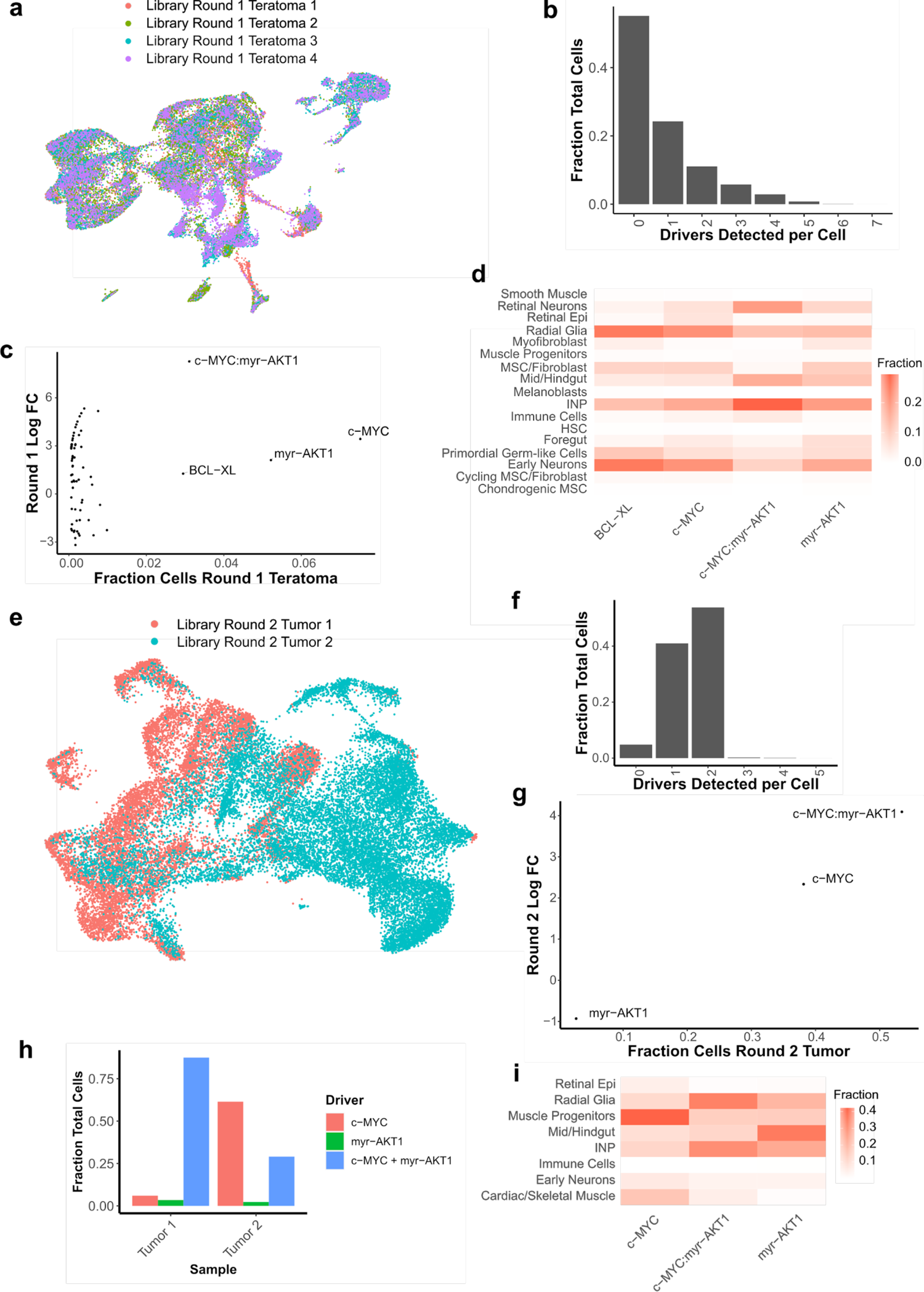
Characterization of driver library screen tumors. (**a**) UMAP visualization of cells by sample from round 1 teratomas formed by driver library transduced hESCs. (**b**) Fraction of cells for each number of drivers detected in driver library round 1 teratomas. (**c**) Scatterplot of log fold change of drivers in round 1 teratomas over pre-injection cells, versus fraction of cells in round 1 teratomas for each driver. Only those drivers detected in at least 25 cells across all 4 round 1 teratomas are plotted. Those drivers detected in at least 1% of round 1 teratoma cells are annotated. (**d**) Heatmap of fraction of cells of each type detected for top detected drivers in round 1 teratomas. (**e**) UMAP visualization of cells by sample from round 1 teratomas formed by driver library transduced hESCs. (**f**) Fraction of cells for each number of drivers detected in driver library round 2 tumors. (**g**) Top enriched hits for each round 2 tumor sample. (**h**) Scatterplot of log fold change of drivers in round 2 tumors over round 1 teratomas, versus fraction of cells in round 2 tumors for each driver. Only those drivers detected in at least 25 cells across both round 2 tumors are plotted. Those drivers detected in at least 1% of round 2 tumor cells are annotated. (**i**) Heatmap of fraction of cells of each type detected for top detected drivers in round 2 tumors.

**Supplementary Figure 3:**
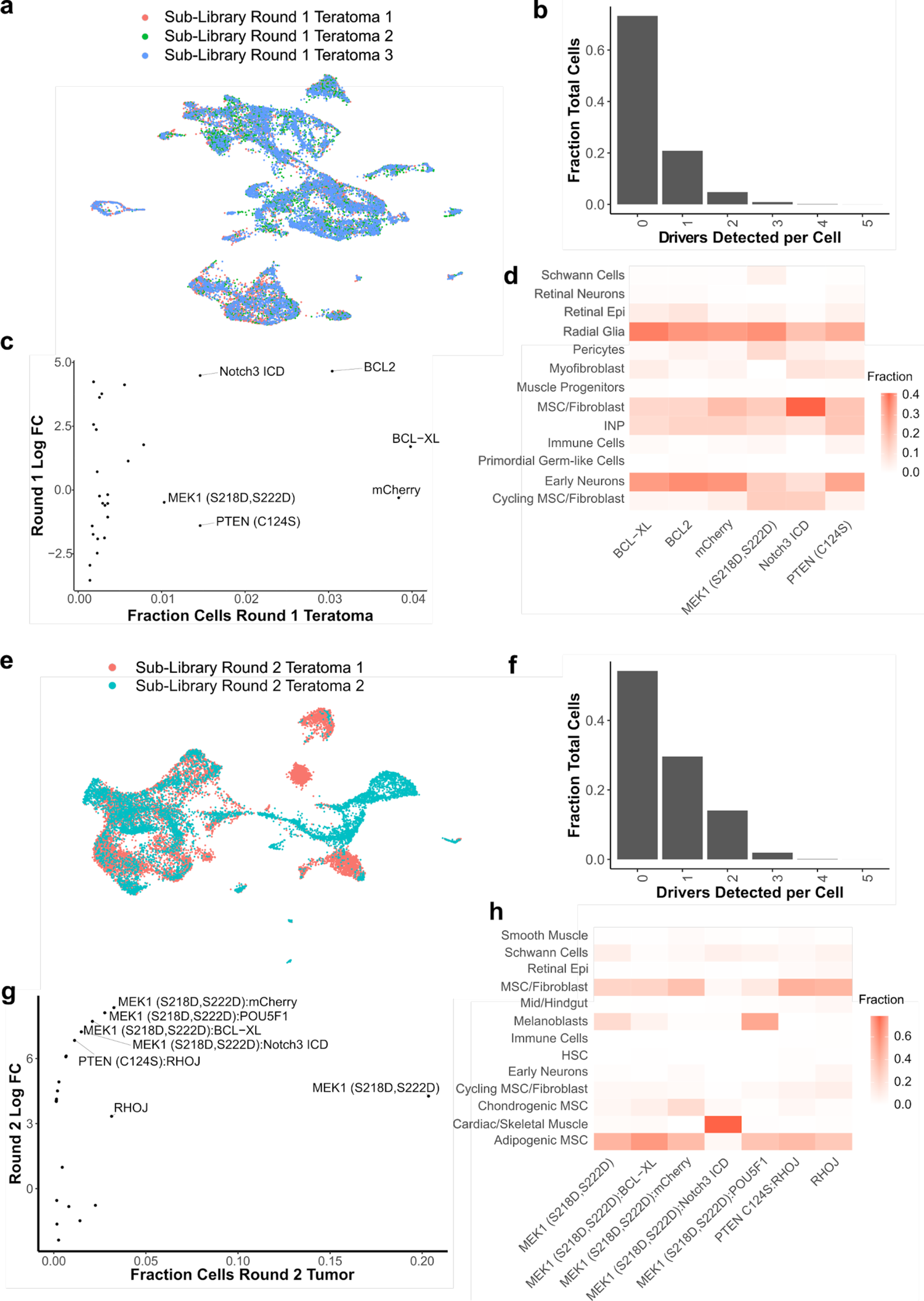
Characterization of driver sub-library screen tumors. (**a**) UMAP visualization of cells by sample from round 1 teratomas formed by driver sub-library transduced hESCs. (**b**) Fraction of cells for each number of drivers detected in driver sub-library round 1 teratomas. (**c**) Scatterplot of log fold change of drivers in round 1 sub-library teratomas over pre-injection cells, versus fraction of cells in round 1 teratomas for each driver. Only those drivers detected in at least 25 cells across all 3 round 1 sub-library teratomas are plotted. Those drivers detected in at least 1% of round 1 teratoma cells are annotated. (**d**) Heatmap of fraction of cells of each type detected for top detected drivers in round 1 teratomas. (**e**) UMAP visualization of cells by sample from round 1 teratomas formed by driver sub-library transduced hESCs. (**f**) Fraction of cells for each number of drivers detected in driver sub-library round 2 tumors. (**g**) Scatterplot of log fold change of drivers in round 2 tumors over round 1 teratomas, versus fraction of cells in round 2 tumors for each driver. Only those drivers detected in at least 25 cells across both round 2 tumors are plotted. Those drivers detected in at least 1% of round 2 tumor cells are annotated. (**h**) Heatmap of fraction of cells of each type detected for top detected drivers in round 2 tumors.

**Supplementary Figure 4:**
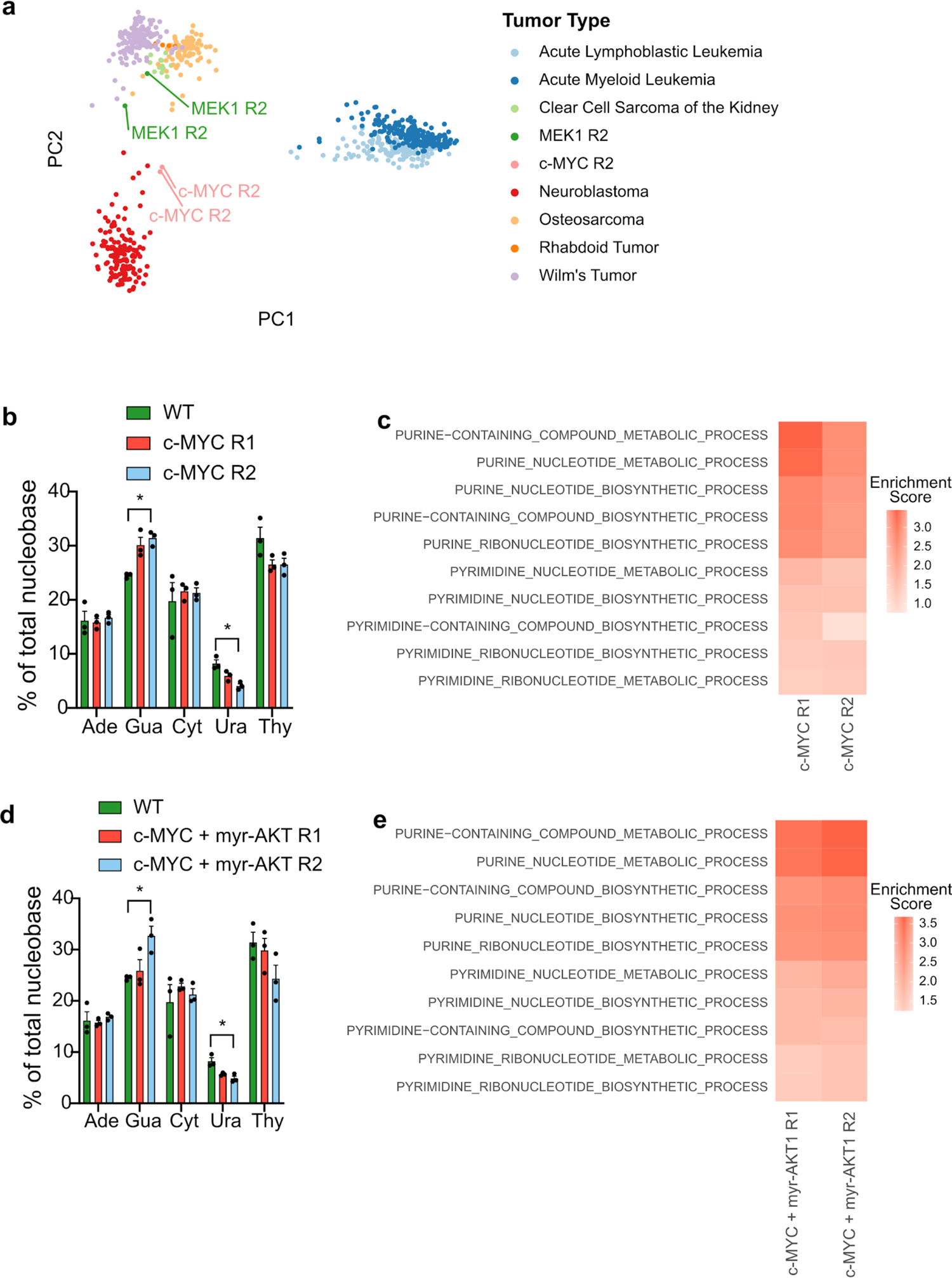
Metabolic Effects of *c-MYC* and *c-MYC* + *myr-AKT1*. (**a**) PC1 vs PC2 plot of *c-MYC* and *MEK1^S^*^218^*^/S222D^* driven tumors with pediatric tumors from the TARGET initiative. (**b**) Nucleobase abundance in Control (Wild Type) teratomas, *c-MYC* driven Round 1 tumors and *c-MYC* driven Round 2 tumors. (**c**) Heatmap of enrichment scores for Gene Ontology genesets related to purine and pyrimidine nucleotide metabolism and synthesis for *c-MYC* driven Round 1 and Round 2 tumors compared to wild type teratomas. (**d**) Nucleobase abundance in Control (Wild Type) teratomas, *c-MYC* + *myr-AKT1* driven Round 1 tumors and *c-MYC* + *myr-AKT1* driven Round 2 tumors. (**e**) Heatmap of enrichment scores for Gene Ontology genesets related to purine and pyrimidine nucleotide metabolism and synthesis for *c-MYC* + *myr-AKT1* driven Round 1 and Round 2 tumors compared to wild type teratomas.

## METHOD DETAILS

### Cell Culture

H1 male hESC cell line was maintained under feeder-free conditions in mTeSR1 medium (Stem Cell Technologies) supplemented with 1% antibiotic-antimycotic (Thermo Fisher Scientific). Prior to passaging, tissue-culture plates were coated with growth factor-reduced Matrigel (Corning) diluted in DMEM/F-12 medium (Thermo Fisher Scientific) and incubated for 30 minutes at 37 ⁰C, 5% CO_2_. Cells were dissociated and passaged using the dissociation reagent Versene (Thermo Fisher Scientific). Prior to lentiviral transduction, hESCs were passaged using Accutase (Innovative Cell Technologies) and plated as dissociated cells to achieve higher transduction efficiency. When passaged with Accutase, cells were plated in mTeSR1 containing Y27632 (10 μM, Tocris). H1 hESCs used started at P30 and were passaged a maximum of 4 passages before injection.

HEK 293T cells were maintained in high glucose DMEM supplemented with 10% fetal bovine serum (FBS) and 1 % antibiotic-antimycotic. HEK 293T cells were passaged using 0.05% Trypsin (Thermo Fisher Scientific).

### Library Preparation

The lentiviral backbone plasmid, compatible with detection in scRNA-seq was constructed as previously described^1^ containing the EF1α promoter, mCherry transgene flanked by BamHI restriction sites, followed by a P2A peptide and hygromycin resistance enzyme gene immediately downstream. Each driver ORF in the library was individually inserted in place of the mCherry transgene. A barcode sequence was introduced to allow for identification of the ectopically expressed transcription factor. The backbone plasmid was digested with HpaI, and a pool of 20 bp long barcodes with flanking sequences compatible with the HpaI site, was inserted immediately downstream of the hygromycin resistance gene by Gibson assembly. The vector was constructed such that the barcodes were located only 200 bp upstream of the 3’-LTR region. This design enabled the barcodes to be transcribed near the poly-adenylation tail of the transcripts and a high fraction of barcodes to be captured during sample processing for scRNA-seq.

To create the driver ORF library, individual drivers were PCR amplified out of the Cancer Pathways kit (Addgene #1000000072)^21^, individual plasmids (Addgene #9053, #85140, #82262, #82297, #82175, #61852, #39872, #23776, #23688, #10745, #23231)^84–89^, a human cDNA pool (Promega Corporation), or obtained as synthesized double-stranded DNA fragments (gBlocks, IDT Inc) with flanking sequences compatible with the BamHI restriction sites. The barcoded lentiviral backbone was digested with BamHI HF (New England Biolabs) at 37 ⁰C for 3 hours in a reaction consisting of: lentiviral backbone, 4 μg, CutSmart buffer, 5 μl, BamHI, 0.625 μl, H_2_0 up to 50 μl. After digestion, the vector was purified using a QIAquick PCR Purification Kit (Qiagen). Each transcription factor vector was then individually assembled via Gibson assembly. The Gibson assembly reactions were set up as follows: 100 ng digested lentiviral backbone, 3:10 molar ratio of transcription factor insert, 2X Gibson assembly master mix (New England Biolabs), H_2_0 up to 20 μl. After incubation at 50 °C for 1 h, the product was transformed into One Shot Stbl3 chemically competent Escherichia coli (Invitrogen). A fraction (150 µL) of cultures was spread on carbenicillin (50 µg/ml) LB plates and incubated overnight at 37 ⁰C. Individual colonies were picked, introduced into 5 ml of carbenicillin (50 µg/ml) LB medium and incubated overnight in a shaker at 37 ⁰C. The plasmid DNA was then extracted with a QIAprep Spin Miniprep Kit (Qiagen), and Sanger sequenced to verify correct assembly of the vector and to extract barcode sequences. One overexpression vector was created for each ORF, thus a single unique barcode was associated with each ORF.

### Viral Production

HEK 293T cells were maintained in high glucose DMEM supplemented with 10% fetal bovine serum (FBS).

To prepare lentivirus for the full library, to avoid barcode shuffling^49–52^ lentivirus for each ORF was packaged independently. Cells were seeded in a 6-well plate 1 day prior to transfection, such that they were 60-70% confluent at the time of transfection. For each well of a 6-well plate, 2.25 μl of Lipofectamine 2000 (Life Technologies) was added to 125 μl of Opti-MEM (Life Technologies). Separately 187.5 ng of pMD2.G (Addgene #12259), 750 ng of pCMV delta R8.2 (Addgene #12263) and 562.5 ng of an individual vector was added to 125 μl of Opti-MEM. After 5 minutes of incubation at room temperature, the Lipofectamine 2000 and DNA solutions were mixed and incubated at room temperature for 30 minutes. During the incubation period, medium in each plate was replaced with 2 ml of fresh, pre-warmed medium per well. After the incubation period, the mixture was added dropwise to each well of HEK 293T cells. Supernatant containing the viral particles was harvested after 48 and 72 hours, filtered with 0.45 μm filters (Steriflip, Millipore), and further concentrated using Amicon Ultra-15 centrifugal ultrafilters with a 100,000 NMWL cutoff (Millipore). For library lentiviral production, supernatant of all ORF wells was mixed together and concentrated to a final volume of 600-800 μl, divided into aliquots and frozen at −80 ⁰C. To prepare lentivirus for individual vectors, the production protocol as described above was scaled up to a 15 cm dish.

### Viral Transduction

For viral transduction, on day −1, H1 cells were dissociated to a single cell suspension using Accutase and seeded into Matrigel-coated 6-well plates at a density of 3×10^5^ cells per well in mTeSR containing ROCK inhibitor, Y27632 (10 μM, Tocris). The next day, day 0, cells were approximately 20% confluent. Medium containing Y27632 was replaced with mTeSR1 within 16 hours after plating and cells were allowed to recover for at least 8 hours prior to addition of virus.

Recovered cells were then transduced with lentivirus added to fresh mTeSR containing polybrene (5 μg/ml, Millipore). On day 1, medium was replaced with fresh mTeSR1. Hygromycin (Thermo Fisher Scientific) selection was started from day 2 onward at a selection dose of 50 μg/ml, medium containing hygromycin was replaced daily.

### Animals

Housing, husbandry and all procedures involving animals used in this study were performed in compliance with protocols approved by the University of California San Diego Institutional Animal Care and Use Committee (UCSD IACUC). Mice were group housed (up to 4 animals per cage) on a 12:12 hr light-dark cycle, with free access to food and water in individually ventilated specific pathogen free (SPF) cages. All mice used were healthy and were not involved in any previous procedures nor drug treatment unless indicated otherwise. All studies performed in NOD.Cg-Prkdcscid Il2rgtm1Wjl/SzJ (NSG) mice and maintained in autoclaved cages.

### Teratoma Formation

A subcutaneous injection of 6-8 million PSCs in a slurry of growth factor reduced Matrigel (Corning) and mTeSR medium (1:1) was made in the right flank of anesthetized Rag2^-/-^;γc^-/-^ immunodeficient mice. Weekly monitoring of teratoma growth was made by quantifying approximate elliptical area (mm^2^) with the use of calipers measuring outward width and height.

### Teratoma Processing

Mice were euthanized by slow release of CO_2_ followed by secondary means via cervical dislocation. Tumor area was shaved, sprayed with 70% ethanol, and then extracted via surgical excision using scissors and forceps. Tumor was rinsed with PBS, weighed, and photographed. Tumor was then cut into small pieces in a semi-random fashion and frozen in OCT for sectioning and H&E staining courtesy of the Moore’s Cancer Center Histology Core. Remaining tumor was cut into small pieces 1-2mm in diameter and subjected to standard GentleMACS (Miltenyi) protocols: Human Tumor Dissociation Kit (medium tumor settings) and Red Blood Cell Lysis Kit. From round 1 teratomas, 6-8 million dissociated live cells were resuspended in growth factor reduced Matrigel (Corning) and subcutaneously injected in the right flank of anesthetized Rag2^-/-^;γc^-/-^ immunodeficient mice to form round 2 tumors. For single cell RNA-seq, samples were also processed with Dead Cell Removal Kit (Miltenyi). Single cells were then resuspended in PBS + 0.04% BSA for processing on the 10X Genomics Chromium^90^ platform and downstream sequencing on an Illumina NovaSeq platform. For bulk processing, after red blood cell removal, cells were divided into multiple tubes and pelleted for 5 minutes at 300g. Supernatant was then removed and cells were either directly frozen at −80 ⁰C or resuspended in RNALater (Thermo Fisher Scientific), incubated overnight at 4 ⁰C, RNALater removed and then frozen at −80 ⁰C.

### Barcode Amplification

Barcodes were amplified from cDNA generated by the single cell system, and gDNA from validation tumors and prepared for deep sequencing through a two-step PCR process.

For amplification of barcodes from cDNA, the first step was performed as four separate 50 μl reactions for each sample. 2.5 μl of the cDNA was input per reaction with Kapa Hifi Hotstart ReadyMix (Kapa Biosystems). The PCR primers used were, NEB_EC2H_Barcode_F: GACTGGAGTTCAGACGTGTGCTCTTCCGATCTAGAACTATTTCCTGGCTGTTACGCG and NEBNext Universal PCR Primer for Illumina (New England Biolabs). The thermocycling parameters were 95 °C for 3 min; 20-26 cycles of (98 °C for 20 s; 65 °C for 15 s; and 72 °C for 30 s); and a final extension of 72 °C for 5 min. The numbers of cycles were tested to ensure that they fell within the linear phase of amplification. Amplicons (∼500 bp) of 4 reactions for each sample were pooled, size-selected and purified with Agencourt AMPure XP beads at a 0.8 ratio. The second step of PCR was performed with two separate 50 μl reactions with 50 ng of first step purified PCR product per reaction. NEBNext Multiplex Oligos for Illumina (Dual Index Primers) were used to attach Illumina adapters and indices to the samples. The thermocycling parameters were: 95 °C for 3 min; 6 cycles of (98 °C for 20 s; 65 °C for 15 s; 72 °C for 30 s); and 72 °C for 5 min. The amplicons from these two reactions for each sample were pooled, size-selected and purified with Agencourt AMPure XP beads at an 0.8 ratio. The purified second-step PCR library was quantified by Qubit dsDNA HS assay (Thermo Fisher Scientific) and used for downstream sequencing on an Illumina HiSeq platform.

For amplification of barcodes from genomic DNA, genomic DNA was extracted from stored cell pellets with a DNeasy Blood and Tissue Kit (Qiagen). The first step PCR was performed as two separate 50 μl reactions for each sample. 2 μg of genomic DNA was input per reaction with Kapa Hifi Hotstart ReadyMix. The PCR primers used were, EC2H_gDNA_Barcode_F: ACACTCTTTCCCTACACGACGCTCTTCCGATCTACTGTCGGGCGTACACAAATC and EC2H_gDNA_Barcode_R: GACTGGAGTTCAGACGTGTGCTCTTCCGATCTCACTGTTTAACAAGCCCGTCAGTAG. The thermocycling parameters were: 95 °C for 3 min; 24-32 cycles of (98 °C for 20 s; 60 °C for 15 s; and 72 °C for 30 s); and a final extension of 72 °C for 5 min. The numbers of cycles were tested to ensure that they fell within the linear phase of amplification. Amplicons (200 bp) of the two reactions for each sample were pooled, size-selected with Agencourt AMPure XP beads (Beckman Coulter, Inc.) at a ratio of 1.6. The second step of PCR was performed as two separate 50 μl reactions with 25 ng of first step purified PCR product per reaction. NEBNext Multiplex Oligos for Illumina (Dual Index Primers) were used to attach Illumina adapters and indices to the samples. The thermocycling parameters were: 95 °C for 3 min; 6-8 cycles of (98 °C for 20 s; 65 °C for 20 s; 72 °C for 30 s); and 72 °C for 2 min. The amplicons from these two reactions for each sample were pooled, size-selected with Agencourt AMPure XP beads at a ratio of 1.6. The purified second-step PCR library was quantified by Qubit dsDNA HS assay (Thermo Fisher Scientific) and used for downstream sequencing on an Illumina NovaSeq platform.

### Single cell RNA-seq Processing

Fastq files were aligned to a combined hg19 and mm10 reference and expression matrices generated using the count command in cellranger v3.0.1 (10X Genomics). cellranger commands were run using default settings.

### Data Integration and Clustering

Data integration was performed on counts matrices from the following samples: 4 Round 1 teratomas perturbed with the full library, 2 Round 2 teratomas perturbed with the full library, 3 Round 1 teratomas perturbed with a library without *c-MYC* or *myr-AKT1*, and 2 Round 2 teratomas perturbed with a library without *c-MYC* or *myr-AKT1*. Integration was done using the Seurat v3 pipeline^53^. Expression matrices were filtered to remove any cells expressing less than 200 genes or expressing greater than 20% mitochondrial genes, as well as to remove any genes that are expressed in less than 0.1% of cells. DoubletFinder^91^ was used to detect predicted doublets, and these were removed for downstream analysis. The expression matrix was then normalized for total counts, log transformed and scaled by a factor of 10,000 for each sample, and the top 4000 most variable genes were identified. We then used Seurat to find anchor cells and integrated all data sets, obtaining a batch-corrected expression matrix for subsequent processing. This expression matrix was then scaled, and nUMI as well as mitochondrial gene fraction was regressed out. Principal component analysis (PCA) was performed on this matrix and 22-30 PCs were identified as significant using an elbow plot. The significant PCs were then used to generate a k Nearest Neighbors (kNN) graph with k=10. The kNN graph was then used to generate a shared Nearest Neighbors (sNN) graph followed by modularity optimization to find clusters with a resolution parameter of 0.8.

To calculate change in the cell type abundance, the number of cells of each type was summarized for each wild type and each driver library sample. This table was input to edgeR^92, 93^, which was then used to determine log_2_ fold change, p-value and false discovery rate (FDR).

### Barcode Assignment

To assign one or more barcodes to each cell, we used a previously described method ^49^. Briefly, we extracted each barcode by identifying its flanking sequences, resulting in reads that contain cell, UMI, and barcode tags. To remove potential chimeric reads, we used a two-step filtering process. First, we only kept barcodes that made up at least 0.5% of the total amount of reads for each cell. We then counted the number of UMIs and reads for each plasmid barcode within each cell, and only assigned that cell any barcode that contained at least 10% of the cell’s read and UMI counts.

### Bulk RNA Extraction and RNA-seq library preparation

RNA was extracted from cells using the RNeasy Mini Kit (Qiagen) as per the manufacturer’s instructions. The quality and concentration of the RNA samples was measured using a spectrophotometer (Nanodrop 2000, Thermo Fisher Scientific).

Bulk RNA-seq libraries were prepared from 1000 ng of RNA using the NEBNext Ultra RNA Library Prep kit for Illumina or the NEBNext Ultra II Directional RNA Library Prep Kit for Illumina (New England Biolabs) as per the manufacturer’s instructions. Libraries were sequenced on an Illumina NovaSeq platform.

### Bulk RNA-Seq Analysis

We mapped the bulk RNA-Seq fastq files to GRCh38 and quantified read counts mapping to each gene’s exon using Ensembl v99 and STAR aligner^94^. The genes by counts matrix was then processed via DESeq2^95^ to normalize counts and estimate differential expression. Log fold change in gene expression of *c-MYC* and *c-MYC* + *myr-AKT1* driven tumors compared to wild type teratomas was used as input for geneset enrichment analysis.

Data from the TARGET initiative was obtained from the UCSC Xena browser. TARGET data as well as experimental tumor samples were processed using edgeR^92^ and limma^96^ to filter and normalize the data and to remove heteroscedascity. The 2000 most highly variable genes across the TARGET tumors were then selected for performing principal component analysis.

### Barcode Counting

Barcodes amplified and sequenced from genomic DNA were aligned using Bowtie2^97, 98^, and then counted using MAGeCK^99^.

### Gas chromatograph-Mass spectrometry (GC-MS) sample preparation and analysis

Metabolites were extracted, analyzed, and quantified, as previously described in detail^100^. Frozen tissue was pulverized using a cellcrusher and ten to fifteen mg of frozen tissue were then homogenized with a ball mill (Retsch Mixer Mill MM 400) at 30 Hz for 5 minutes in 500 µL −20°C methanol and 200 µL of ice-cold MiliQ water. The mixture was then transferred into a 2 mL Eppendorf tube containing in 500 µL of chloroform, vortexed for 5 min followed by centrifugation with 16 000 x *g* for 5 min at 4 °C. To determine the abundances of nucleobases the interface was centrifuged with 1mL −20 °C methanol at 16,000×*g* for 5 min at 4 °C, the pellet was hydrolyzed with 1 mL 6N HCL for 2h at 80°C, centrifuged at 16,000×*g* for 5 min and the supernatant was dried at 60 °C under airflow.

Metabolite derivatization was performed using a Gerstel MPS. Dried metabolites were dissolved in 15 μl of 2% (w/v) methoxyamine hydrochloride (Thermo Scientific) in pyridine and incubated for 60 min at 45°C. An equal volume of 2,2,2-trifluoro-N-methyl-N-trimethylsilyl-acetamide (MSTFA) (nucleobases) was added and incubated further for 30 min at 45 °C. Derivatized samples were analyzed by GC-MS using a DB-35MS column (30 m x 0.25 mm i.d. x 0.25 µm, Agilent J&W Scientific) installed in an Agilent 7890A gas chromatograph (GC) interfaced with an Agilent 5975C mass spectrometer (MS) operating under electron impact ionization at 70 eV. The MS source was held at 230 °C and the quadrupole at 150 °C and helium was used as a carrier gas at a flow rate of 1 mL/min. The GC oven temperature was held at 80 °C for 6 min, increased to 300 °C at 6 °C/min and after 10 min increased to 325 °C at 10 °C/min for 4 min.

## REFERENCES

1. Merlo, L. M. F., Pepper, J. W., Reid, B. J. & Maley, C. C. Cancer as an evolutionary and ecological process. Nature Reviews Cancer vol. 6 924–935 (2006).

2. Greaves, M. & Maley, C. C. Clonal evolution in cancer. Nature vol. 481 306–313 (2012).

3. Hagis, K. M., Cichowski, K. & Elledge, S. J. Tissue-specificity in cancer: The rule, not the exception. Science 363, 1150–1151 (2019).

4. Schneider, G., Schmidt-Supprian, M., Rad, R. & Saur, D. Tissue-specific tumorigenesis: Context matters. Nature Reviews Cancer vol. 17 239–253 (2017).

5. Sottoriva, A. et al. A Big Bang model of human colorectal tumor growth. Nat. Genet. 47, 209– 216 (2015).

6. Beroukhim, R. et al. The landscape of somatic copy-number alteration across human cancers. Nature 463, 899–905 (2010).

7. Bailey, M. H. et al. Comprehensive Characterization of Cancer Driver Genes and Mutations. Cell 173, 371–385.e18 (2018).

8. Alexandrov, L. B. et al. Signatures of mutational processes in human cancer. Nature 500, 415–421 (2013).

9. Forbes, S. A. et al. COSMIC: Somatic cancer genetics at high-resolution. Nucleic Acids Res. 45, D777–D783 (2017).

10. Zack, T. I. et al. Pan-cancer patterns of somatic copy number alteration. Nat. Genet. 45, 1134–1140 (2013).

11. Sanchez-Vega, F. et al. Oncogenic Signaling Pathways in The Cancer Genome Atlas. Cell 173, 321–337.e10 (2018).

12. Etzioni, R. et al. The case for early detection. Nature Reviews Cancer vol. 3 243–252 (2003).

13. Balani, S., Nguyen, L. V. & Eaves, C. J. Modeling the process of human tumorigenesis. Nature Communications vol. 8 1–10 (2017).

14. Richmond, A. & Yingjun, S. Mouse xenograft models vs GEM models for human cancer therapeutics. DMM Disease Models and Mechanisms vol. 1 78–82 (2008).

15. Gould, S. E., Junttila, M. R. & De Sauvage, F. J. Translational value of mouse models in oncology drug development. Nature Medicine vol. 21 431–439 (2015).

16. Cheon, D.-J. & Orsulic, S. Mouse Models of Cancer. Annu. Rev. Pathol.: Mech. Dis. 6, 95– 119 (2011).

17. Rangarajan, A. & Weinberg, R. A. Comparative biology of mouse versus human cells: Modelling human cancer in mice. Nature Reviews Cancer vol. 3 952–959 (2003).

18. Fischer, M. Mice Are Not Humans: The Case of p53. Trends in Cancer vol. 7 12–14 (2021).

19. Wilding, J. L. & Bodmer, W. F. Cancer cell lines for drug discovery and development. Cancer Res. 74, 2377–2384 (2014).

20. Gillet, J.-P., Varma, S. & Gottesman, M. M. The clinical relevance of cancer cell lines. J. Natl. Cancer Inst. 105, 452–458 (2013).

21. Martz, C. A. et al. Systematic identification of signaling pathways with potential to confer anticancer drug resistance. Sci. Signal. 7, 1–14 (2014).

22. Hahn, W. C. et al. Creation of human tumour cells with defined genetic elements. Nature 400, 464–468 (1999).

23. Rangarajan, A., Hong, S. J., Gifford, A. & Weinberg, R. A. Species- and cell type-specific requirements for cellular transformation. Cancer Cell 6, 171–183 (2004).

24. Elenbaas, B. et al. Human breast cancer cells generated by oncogenic transformation of primary mammary epithelial cells. Genes and Development 15, 50–65 (2001).

25. Daley, G. Q., Mclaughlin, J., Witte, O. N. & Baltimore, D. The CML-specific P210 bcr/abl protein, unlike v-abl, does not transform NIH/3T3 fibroblasts. Science 237, 532–535 (1987).

26. Clark, R. et al. Transformation of Human Mammary Epithelial Cells by Oncogenic Retroviruses. vol. 48 4689–4694 (1988).

27. Sasaki, R. et al. Oncogenic transformation of human ovarian surface epithelial cells with defined cellular oncogenes. Carcinogenesis 30, 423–431 (2009).

28. Boehm, J. S., Hession, M. T., Bulmer, S. E. & Hahn, W. C. Transformation of Human and Murine Fibroblasts without Viral Oncoproteins. Mol. Cell. Biol. 25, 6464–6474 (2005).

29. Geder, L., Lausch, R., O’Neill, F. & Rapp, F. Oncogenic transformation of human embryo lung cells by human cytomegalovirus. Science 192, 1134–1137 (1976).

30. Park, J. W. et al. Reprogramming normal human epithelial tissues to a common, lethal neuroendocrine cancer lineage. Science 362, 91–95 (2018).

31. Sack, L. M. et al. Profound Tissue Specificity in Proliferation Control Underlies Cancer Drivers and Aneuploidy Patterns. Cell 0, 1–16 (2018).

32. Puisieux, A., Pommier, R. M., Morel, A. P. & Lavial, F. Cellular Pliancy and the Multistep Process of Tumorigenesis. Cancer Cell vol. 33 164–172 (2018).

33. Smith, R. C. & Tabar, V. Constructing and Deconstructing Cancers using Human Pluripotent Stem Cells and Organoids. Cell Stem Cell vol. 24 12–24 (2019).

34. Drost, J. et al. Sequential cancer mutations in cultured human intestinal stem cells. Nature 521, 43–47 (2015).

35. Matano, M. et al. Modeling colorectal cancer using CRISPR-Cas9-mediated engineering of human intestinal organoids. Nat. Med. 21, 256–262 (2015).

36. Bian, S. et al. Genetically engineered cerebral organoids model brain tumor formation. Nat. Methods 15, 631–639 (2018).

37. Drost, J. et al. Use of CRISPR-modified human stem cell organoids to study the origin of mutational signatures in cancer. Science 358, 234–238 (2017).

38. Fumagalli, A. et al. Genetic dissection of colorectal cancer progression by orthotopic transplantation of engineered cancer organoids. Proc. Natl. Acad. Sci. U. S. A. 114, E2357– E2364 (2017).

39. Rosenbluth, J. M. et al. Organoid cultures from normal and cancer-prone human breast tissues preserve complex epithelial lineages. Nat. Commun. 11, 1711 (2020).

40. Li, X. et al. Oncogenic transformation of diverse gastrointestinal tissues in primary organoid culture. Nat. Med. 20, 769–777 (2014).

41. Lannagan, T. R. M. et al. Genetic editing of colonic organoids provides a molecularly distinct and orthotopic preclinical model of serrated carcinogenesis. Gut 68, 684–692 (2019).

42. Hanahan, D. & Weinberg, R. A. Hallmarks of cancer: The next generation. Cell 144, 646– 674 (2011).

43. Koga, T. et al. Longitudinal assessment of tumor development using cancer avatars derived from genetically engineered pluripotent stem cells. Nat. Commun. 11, 550 (2020).

44. Duan, S. et al. PTEN deficiency reprogrammes human neural stem cells towards a glioblastoma stem cell-like phenotype. Nat. Commun. 6, 10068 (2015).

45. Pei, Y. et al. An Animal Model of MYC-Driven Medulloblastoma. Cancer Cell 21, 155–167 (2012).

46. Pei, Y. et al. HDAC and PI3K Antagonists Cooperate to Inhibit Growth of MYC-Driven Medulloblastoma. Cancer Cell 29, 311–323 (2016).

47. Lensch, M. W., Schlaeger, T. M., Zon, L. I. & Daley, G. Q. Teratoma Formation Assays with Human Embryonic Stem Cells: A Rationale for One Type of Human-Animal Chimera. Cell Stem Cell vol. 1 253–258 (2007).

48. McDonald, D. et al. Defining the Teratoma as a Model for Multi-lineage Human Development. Cell 183, 1402–1419.e18 (2020).

49. Parekh, U. et al. Mapping Cellular Reprogramming via Pooled Overexpression Screens with Paired Fitness and Single-Cell RNA-Sequencing Readout. Cell systems 7, 548–555.e8 (2018).

50. Sack, L. M., Davoli, T., Xu, Q., Li, M. Z. & Elledge, S. J. Sources of Error in Mammalian Genetic Screens. G3 6, 2781–2790 (2016).

51. Hill, A. J. et al. On the design of CRISPR-based single-cell molecular screens. Nat. Methods 15, 271–274 (2018).

52. Xie, S., Cooley, A., Armendariz, D., Zhou, P. & Hon, G. C. Frequent sgRNA-barcode recombination in single-cell perturbation assays. PLoS One 13, e0198635 (2018).

53. Butler, A., Hoffman, P., Smibert, P., Papalexi, E. & Satija, R. Integrating single-cell transcriptomic data across different conditions, technologies, and species. Nat. Biotechnol. 36, 411–420 (2018).

54. Fan, J., Slowikowski, K. & Zhang, F. Single-cell transcriptomics in cancer: computational challenges and opportunities. Exp. Mol. Med. 52, 1452–1465 (2020).

55. Yuan, J. et al. Single-cell transcriptome analysis of lineage diversity in high-grade glioma. Genome Med. 10, 57 (2018).

56. Couturier, C. P. et al. Single-cell RNA-seq reveals that glioblastoma recapitulates a normal neurodevelopmental hierarchy. Nat. Commun. 11, 3406 (2020).

57. Vladoiu, M. C. et al. Childhood cerebellar tumours mirror conserved fetal transcriptional programs. Nature 572, 67–73 (2019).

58. Hovestadt, V. et al. Resolving medulloblastoma cellular architecture by single-cell genomics. Nature 572, 74–79 (2019).

59. Ho, B. et al. Molecular subgrouping of atypical teratoid/rhabdoid tumors—a reinvestigation and current consensus. Neuro. Oncol. 22, 613–624 (2020).

60. Alimova, I. et al. Inhibition of *MYC* attenuates tumor cell self-renewal and promotes senescence in SMARCB1-deficient Group 2 atypical teratoid rhabdoid tumors to suppress tumor growth *in vivo*. International Journal of Cancer 144, 1983–1995 (2019).

61. Gravina, G. L. et al. C-Myc Sustains Transformed Phenotype and Promotes Radioresistance of Embryonal Rhabdomyosarcoma Cell Lines. Radiat. Res. 185, 411–422 (2016).

62. Zhang, J. et al. C-Myc promotes tumor proliferation and anti-apoptosis by repressing p21 in rhabdomyosarcomas. Mol. Med. Rep. 16, 4089–4094 (2017).

63. Kouraklis, G., Triche, T. J., Tsokos, M. & Wesley, R. D. Myc oncogene expression and nude mouse tumorigenicity and metastasis formation are higher in alveolar than embryonal rhabdomyosarcoma cell lines. Pediatr. Res. 45, 552–558 (1999).

64. Bouchard, C., Marquardt, J., Brás, A., Medema, R. H. & Eilers, M. Myc-induced proliferation and transformation require Akt-mediated phosphorylation of FoxO proteins. EMBO J. 23, 2830–2840 (2004).

65. Adameyko, I. et al. Schwann Cell Precursors from Nerve Innervation Are a Cellular Origin of Melanocytes in Skin. Cell 139, 366–379 (2009).

66. Brunet, A., Pagès, G. & Pouysségur, J. Constitutively active mutants of MAP kinase kinase (MEK1) induce growth factor-relaxation and oncogenicity when expressed in fibroblasts. Oncogene 9, 3379–3387 (1994).

67. Mansour, S. J. et al. Transformation of mammalian cells by constitutively active MAP kinase kinase. Science 265, 966–970 (1994).

68. Cowley, S., Paterson, H., Kemp, P. & Marshall, C. J. Activation of MAP kinase kinase is necessary and sufficient for PC12 differentiation and for transformation of NIH 3T3 cells. Cell 77, 841–852 (1994).

69. Jin, S. J., Jin, M. Z., Xia, B. R. & Jin, W. L. Long Non-coding RNA DANCR as an Emerging Therapeutic Target in Human Cancers. Frontiers in Oncology vol. 9 1225 (2019).

70. Zhang, J., Tao, Z. & Wang, Y. Long non-coding RNA DANCR regulates the proliferation and osteogenic differentiation of human bone-derived marrow mesenchymal stem cells via the p38 MAPK pathway. Int. J. Mol. Med. 41, 213–219 (2018).

71. Stine, Z. E., Walton, Z. E., Altman, B. J., Hsieh, A. L. & Dang, C. V. MYC, Metabolism, and Cancer. Cancer Discov. 5, 1024–1039 (2015).

72. Hoxhaj, G. & Manning, B. D. The PI3K-AKT network at the interface of oncogenic signalling and cancer metabolism. Nat. Rev. Cancer 20, 74–88 (2020).

73. Liu, Y.-C. et al. Global regulation of nucleotide biosynthetic genes by c-Myc. PLoS One 3, e2722 (2008).

74. Wang, X. et al. Purine synthesis promotes maintenance of brain tumor initiating cells in glioma. Nat. Neurosci. 20, 661–673 (2017).

75. Barfeld, S. J. et al. Myc-dependent purine biosynthesis affects nucleolar stress and therapy response in prostate cancer. Oncotarget 6, 12587–12602 (2015).

76. O’Flanagan, C. H. et al. Dissociation of solid tumor tissues with cold active protease for single-cell RNA-seq minimizes conserved collagenase-associated stress responses. Genome Biol. 20, 210 (2019).

77. Denisenko, E. et al. Systematic assessment of tissue dissociation and storage biases in single-cell and single-nucleus RNA-seq workflows. Genome Biol. 21, 130 (2020).

78. Van Den Brink, S. C. et al. Single-cell sequencing reveals dissociation-induced gene expression in tissue subpopulations. Nature Methods vol. 14 935–936 (2017).

79. Chan, S. S. K. et al. Skeletal Muscle Stem Cells from PSC-Derived Teratomas Have Functional Regenerative Capacity. Cell Stem Cell 23, 74–85.e6 (2018).

80. Walsh, N. C. et al. Humanized Mouse Models of Clinical Disease. Annual Review of Pathology: Mechanisms of Disease vol. 12 187–215 (2017).

81. Rodriques, S. G. et al. Slide-seq: A scalable technology for measuring genome-wide expression at high spatial resolution. Science 363, 1463–1467 (2019).

82. Vickovic, S. et al. High-definition spatial transcriptomics for in situ tissue profiling. Nat. Methods 16, 987–990 (2019).

83. Ståhl, P. L. et al. Visualization and analysis of gene expression in tissue sections by spatial transcriptomics. Science 353, 78–82 (2016).

84. Hayer, A. et al. Engulfed cadherin fingers are polarized junctional structures between collectively migrating endothelial cells. Nat. Cell Biol. 18, 1311–1323 (2016).

85. Kim, E. et al. Systematic Functional Interrogation of Rare Cancer Variants Identifies Oncogenic Alleles. Cancer Discov. 6, 714–726 (2016).

86. Hagting, A., Karlsson, C., Clute, P., Jackman, M. & Pines, J. MPF localization is controlled by nuclear export. EMBO J. 17, 4127–4138 (1998).

87. Johannessen, C. M. et al. COT drives resistance to RAF inhibition through MAP kinase pathway reactivation. Nature 468, 968–972 (2010).

88. Ramaswamy, S. et al. Regulation of G1 progression by the PTEN tumor suppressor protein is linked to inhibition of the phosphatidylinositol 3-kinase/Akt pathway. Proc. Natl. Acad. Sci. U. S. A. 96, 2110–2115 (1999).

89. Roberts, P. J. et al. Rho Family GTPase modification and dependence on CAAX motif-signaled posttranslational modification. J. Biol. Chem. 283, 25150–25163 (2008).

90. Zheng, G. X. Y. et al. Massively parallel digital transcriptional profiling of single cells. Nat. Commun. 8, 1–12 (2017).

91. McGinnis, C. S., Murrow, L. M. & Gartner, Z. J. DoubletFinder: Doublet Detection in Single-Cell RNA Sequencing Data Using Artificial Nearest Neighbors. Cell Systems 8, 329–337.e4 (2019).

92. Robinson, M. D., McCarthy, D. J. & Smyth, G. K. edgeR: a Bioconductor package for differential expression analysis of digital gene expression data. Bioinformatics 26, 139–140 (2010).

93. Ritchie, M. E. et al. edgeR: A versatile tool for the analysis of shRNA-seq and CRISPR-Cas9 genetic screens. F1000Res. 3, (2014).

94. Dobin, A. et al. STAR: Ultrafast universal RNA-seq aligner. Bioinformatics 29, 15–21 (2013).

95. Love, M. I., Huber, W. & Anders, S. Moderated estimation of fold change and dispersion for RNA-seq data with DESeq2. Genome Biol. 15, 550 (2014).

96. Ritchie, M. E. et al. limma powers differential expression analyses for RNA-sequencing and microarray studies. Nucleic Acids Res. 43, e47 (2015).

97. Langmead, B. & Salzberg, S. L. Fast gapped-read alignment with Bowtie 2. Nat. Methods 9, 357–359 (2012).

98. Langmead, B., Trapnell, C., Pop, M. & Salzberg, S. L. Ultrafast and memory-efficient alignment of short DNA sequences to the human genome. Genome Biol. 10, R25 (2009).

99. Li, W. et al. MAGeCK enables robust identification of essential genes from genome-scale CRISPR/Cas9 knockout screens. Genome Biol. 15, 554 (2014).

100. Cordes, T. & Metallo, C. M. Quantifying Intermediary Metabolism and Lipogenesis in Cultured Mammalian Cells Using Stable Isotope Tracing and Mass Spectrometry. Methods Mol. Biol. 1978, 219–241 (2019).

